# The miR-424/503 cluster modulates Wnt/β-catenin signaling in the mammary epithelium by regulating the expression of the LRP6 co-receptor

**DOI:** 10.1101/2021.05.06.442934

**Authors:** Erin A. Nekritz, Ruth Rodriguez-Barrueco, Koon-Kiu Yan, Meredith L. Davis, Rachel L. Werner, Laura Devis-Jauregui, Partha Mukhopadhyay, Jiyang Yu, David Llobet-Navas, Jose Silva

## Abstract

During the female lifetime, the enlargement of the epithelial compartment dictated by the ovarian cycles is supported by a transient increase in the MaSC population. Notably, activation of Wnt/β-catenin signaling is an important trigger for MaSC expansion. Here, we report that the miR-424/503 cluster is a novel modulator of canonical Wnt-signaling in the mammary epithelium that exerts its function by targeting the LRP6 co-receptor. Additionally, we show that the loss of this microRNA cluster is associated with breast cancers possessing high levels of Wnt/β-catenin signaling.

## Introduction

The mammary gland is a very dynamic organ that undergoes continuous tissue remodeling during the female lifetime (Macias & Hinck, 2012). Expansion and regression of the mammary epithelium dictated by the ovarian hormones involve alternating cycles of cell death and proliferation. During this process, enlargement of the epithelial compartment is supported by a transient increase in the mammary epithelial stem cell population (MaSC) (Asselin-Labat *et al*, 2010; Joshi *et al*, 2010). These cycles are tightly regulated in time and space to ensure proper organ function and avoid pathological consequences.

Numerous studies have elucidated some of the major signaling events within the mammary epithelium. For instance, TGFβ has been implicated as a negative regulator of proliferation during ductal elongation (Macias & Hinck, 2012). The induction of apoptosis via the Jak/Stat pathway as well as a number of matrix metalloproteases (MMPs) are involved in the remodeling of the mammary epithelium during involution (Watson, 2006a). RANK ligand is critical for lobuloalveologenesis (Fernandez-Valdivia *et al*, 2009). Importantly, canonical Wnt signaling via β-catenin has been shown to be essential for mammary gland development (Alexander et al., 2012). Specifically, paracrine Wnt-signaling mediated by multiple Wnt-ligands induces mammary epithelial expansion through its effect on MaSCs. In these cells, the expression of Wnt co-receptors LRP5 and LRP6 is the limiting factor for canonical Wnt signaling (Alexander *et al*, 2012; Goel *et al*, 2012). Remarkably, deregulation of Wnt/β-catenin signaling in the mammary gland induces pathological consequences, and hyperactivation of this pathway has been found in some aggressive breast cancer subtypes (Xu *et al*, 2020). Despite the above, our understanding of how canonical Wnt signaling is properly regulated to maintain mammary gland homeostasis is far from complete.

Recently, we have investigated the landscape of miRNAs expressed in the mammary epithelium and found that the miR-424(322)/503 is a TGF-β-induced regulator of involution in the mammary epithelium (Llobet-Navas *et al*, 2014a). This cluster expresses two microRNAs, miR-424(miR-322 in mice) and miR-503, that belong to the miR-16 family (Rissland *et al*, 2011). We have also described that this microRNA cluster is commonly lost in breast cancers and that miR-424(322)/503 knock-out mice develop mammary tumors over time that are promoted by pregnancy (Rodriguez-Barrueco *et al*, 2017). Remarkably, we noticed that tumors emerging in miR-424(322)/503^-/-^ mice presented phenotypical characteristics found in tumors with activated canonical Wnt-signaling. Thus, we decided to investigate a potential link between the loss of miR-424/503 and the Wnt/β-catenin pathway. Here, we report that the miR-424/503 cluster is a novel modulator of the canonical Wnt signaling in the mammary epithelium that exert its function by decreasing the expression of the LRP6 co-receptor. Additionally, we also show that the loss of this miR-cluster is associated with breast cancers with hyperactivation of Wnt/β-catenin signaling.

## Results and Discussion

Commonly, mammary tumors emerging in transgenic mice expressing potent oncogenes present aggressive anaplastic and metaplastic characteristics (Li *et al*, 2003). In the cases where some differentiated characteristics are present, these tumors tend to express defined markers of luminal or basal cancers but rarely both (Pfefferle *et al*, 2013). In contrast to this, we observed that tumors emerging in miR-424/503^-/-^ mice were moderately differentiated and comprised of ducts with layers of epithelial cells showing a high nucleus-to-cytoplasm ratio and extensive fibrosis (Fig. 1A). Additionally, immunohistochemistry staining with cytokeratin (CK) markers revealed that these tumors contained a mix of both luminal and basal cell types (Supplemental Fig. S1A). As these characteristics resemble tumors with activated Wnt/β-catenin signaling we decided to investigate the staining of β-catenin as well as CK6, a marker associated with precursor cells that are found in tumors with hyperactivated canonical Wnt-signaling (Li *et al*., 2003). Remarkably, we found strong staining of both markers including accumulation of nuclear β-catenin in miR-424 /503^-/-^ tumors (Fig. 1A and Supplemental Fig. S1B).

**Figure 1.**
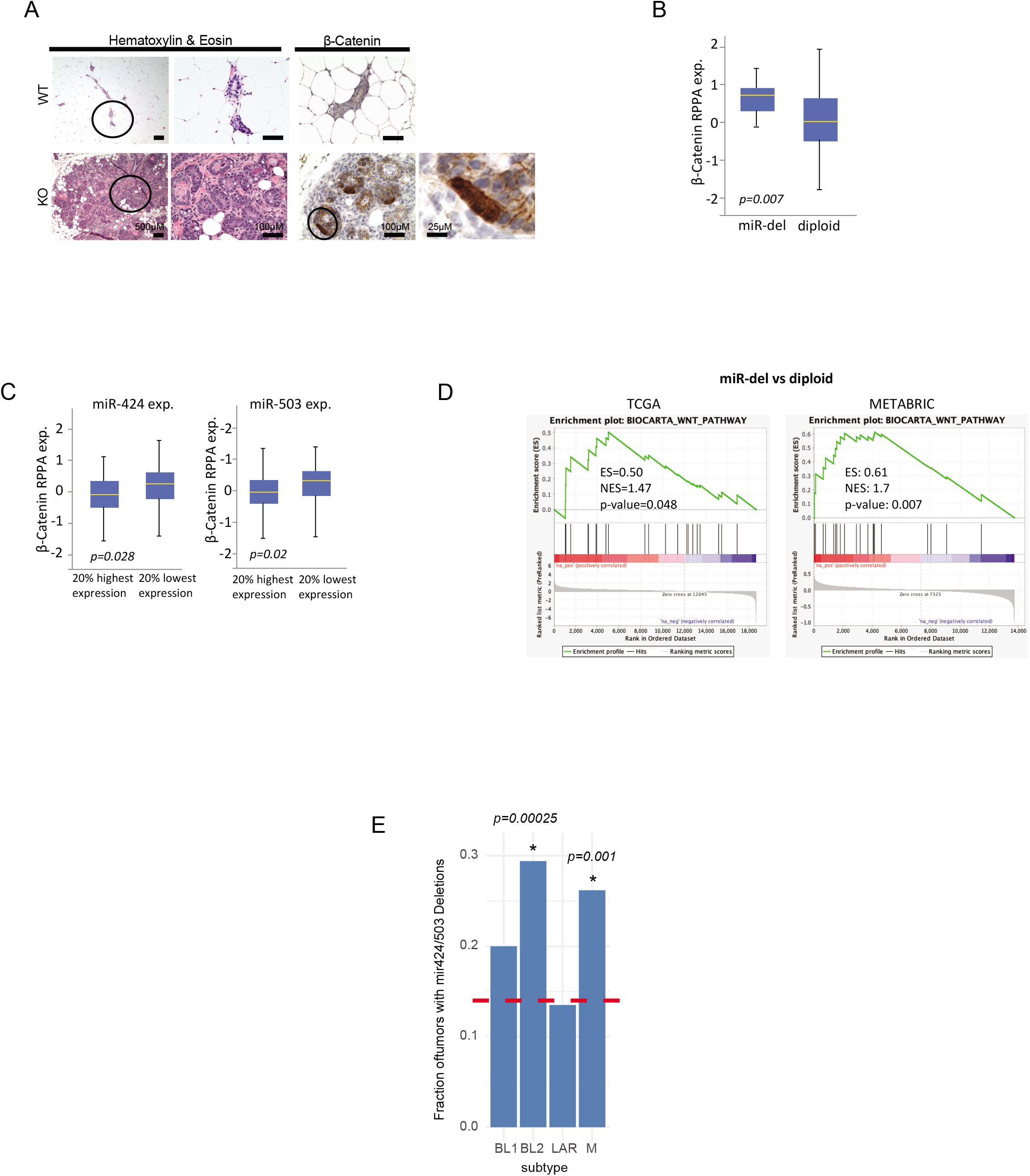
Loss of miR-424/503 is associated with tumors with elevated Wnt/β-catenin activity. A) H&E and immunohistochemistry reveal that β-catenin accumulates in tumors emerging in one-year-old female miR-424(322)/503^-/-^ mice. The panel is a representative image of n=5 independent tumors. B) Bar graphic showing that human breast cancers with deletion of the miR-424/503 locus have higher protein levels of β-catenin (RPPA). P-value is calculated based on one-sided Wilcoxon rank-sum test. C) The bar graph shows that human breast cancers with low expression levels of miR-424 have higher protein levels of β-catenin. P-value is calculated based on one-sided Wilcoxon rank-sum test. D) GSEA indicating that tumors with deletion of the miR-424/503 locus present higher canonical Wnt-signaling activity. Independent analysis for TCGA and METABRIC data sets are shown. E) The graphic shows the enrichment of miR-424/503 deletions in the BL2 and M subtypes of TNBC. The red line shows the overall fraction of miR-424/503 deletions in TCGA samples. The enrichment in subtypes is estimated by the binomial test.

We have previously reported that the miR-424/503 locus is deleted in ∼14% of all breast cancers and that this correlates with lower expression of both miR-424 and miR-503 (Rodriguez-Barrueco *et al*., 2017). Thus, we reasoned that we could investigate the link between the miR-424/503 and canonical Wnt-signaling in human samples by evaluating the accumulation of β-catenin in breast cancers with deletion and/or reduced expression of the miR-cluster. For this, we analyzed the reverse phase protein arrays data (RPPA) that are available for the BRCA-TCGA data set (Cancer Genome Atlas, 2012) and which contains information regarding β-catenin expression at a protein level. In agreement with the observation in mice, both deletion and low expression of miR-424/503 were associated with accumulation of β-catenin (Fig. 1B-1C). Additionally, we performed gene set enrichment analysis to investigate whether this high level of β-catenin impacts the activity of Wnt-signaling. As expected, a significant increase of Wnt-targets was found in tumors deficient in miR-424/503 expression (Fig. 1D). Importantly, this association was confirmed when an independent breast cancer data set containing CNA and expression data for over 2,500 samples (METABRIC (Curtis *et al*, 2012)) was analyzed (Fig 1D). It is well known that a fraction of human breast cancers enriched in triple-negative characteristics have hyperactivation of Wnt-signaling (Pohl *et al*, 2017). However, alterations in signaling components that are commonly found mutated in other tumors (ex. APC, β-catenin) are rare in breast tumors (Sanchez-Vega *et al*, 2018). Genomics analysis has recently classified triple-negative breast cancers (TNBC) in at least four subtypes with different molecular characteristics (Lehmann *et al*, 2011; Lehmann *et al*, 2016). Remarkably, these include two subtypes (∼50% of all TNBCs), basal-like 2 (BL2) and mesenchymal (M), characterized by signatures of epithelial-mesenchymal-transition and hyperactivation of Wnt-signaling (Lehmann *et al*., 2011; Lehmann *et al*., 2016). Thus, we wondered whether deletion of miR-424/503 could be associated with these molecular subtypes. Remarkably, we found a strong association of miR-424/503 deletion with the BL2 and M subtypes (Fig. 1E).

Overall, all the above confirms that deregulation of miR-424/503 influences canonical Wnt-signaling, and that deletion of this microRNA cluster is associated with breast cancers with hyperactivation of Wnt-signaling.

In the mammary gland, Wnt/β-catenin signaling is a critical regulator of epithelial tree homeostasis (Alexander et al., 2012). During regular ovarian cycles and pregnancy, elevated hormonal level promotes secretion of Wnt-ligands in the luminal cells which in turn induces the expansion of MaSCs (Diaz-Guerra *et al*, 2012; Joshi *et al*., 2010). Expansion of MaSCs supports the enlargement of the mammary epithelial compartment. Previously, we have reported that miR-424(322)/503 is expressed at a low level in resting luminal and basal mammary epithelial subpopulations (Llobet-Navas *et al*., 2014a; Llobet-Navas *et al*, 2014b). Based on the new data connecting this cluster with Wnt-signaling we decided to look in more detail its expression in MaSCs. For this, we purified the basal, luminal, and an enriched MaSC subpopulation from 4–6-month-old WT FVB female mice in the estrus and diestrus phase; then, expression of miR-424(322) and miR-503 was assessed by QRT-PCR. In brief, non-epithelial cells were removed from single-cell suspension by incubation with CD31-, CD-45-, Ter119- and BP-1-biotin. The remaining epithelial cells were separated by FACS into two main populations, basal (EpCAM^low^CD49f^hi^) and luminal (EpCAM^high^CD49f^low^) (Smalley *et al*, 2012). Luminal cells were further subdivided in hormone receptor-positive and negative (HR+/-) by Sca1 (Ly6a/e) levels, HR-(EpCAM^high^CD49f^low^Sca-1^-^) and HR+ (EpCAM^high^CD49f^low^Sca-1^+^) (Smalley *et al*., 2012). Finally, a population enriched in MaSCs was purified from the basal compartment by PROCR expression, MaSC EpCAM^low^CD49f^high^PROCR^+^ (Wang *et al*, 2015). Remarkably, the expression of miR-424(322) was found to be up to 30-fold higher in MaSCs than in any other subpopulation during the estrus phase. However, during the expansion phase of the epithelium (diestrus), its expression was strongly reduced (Fig. 2A).

**Figure 2.**
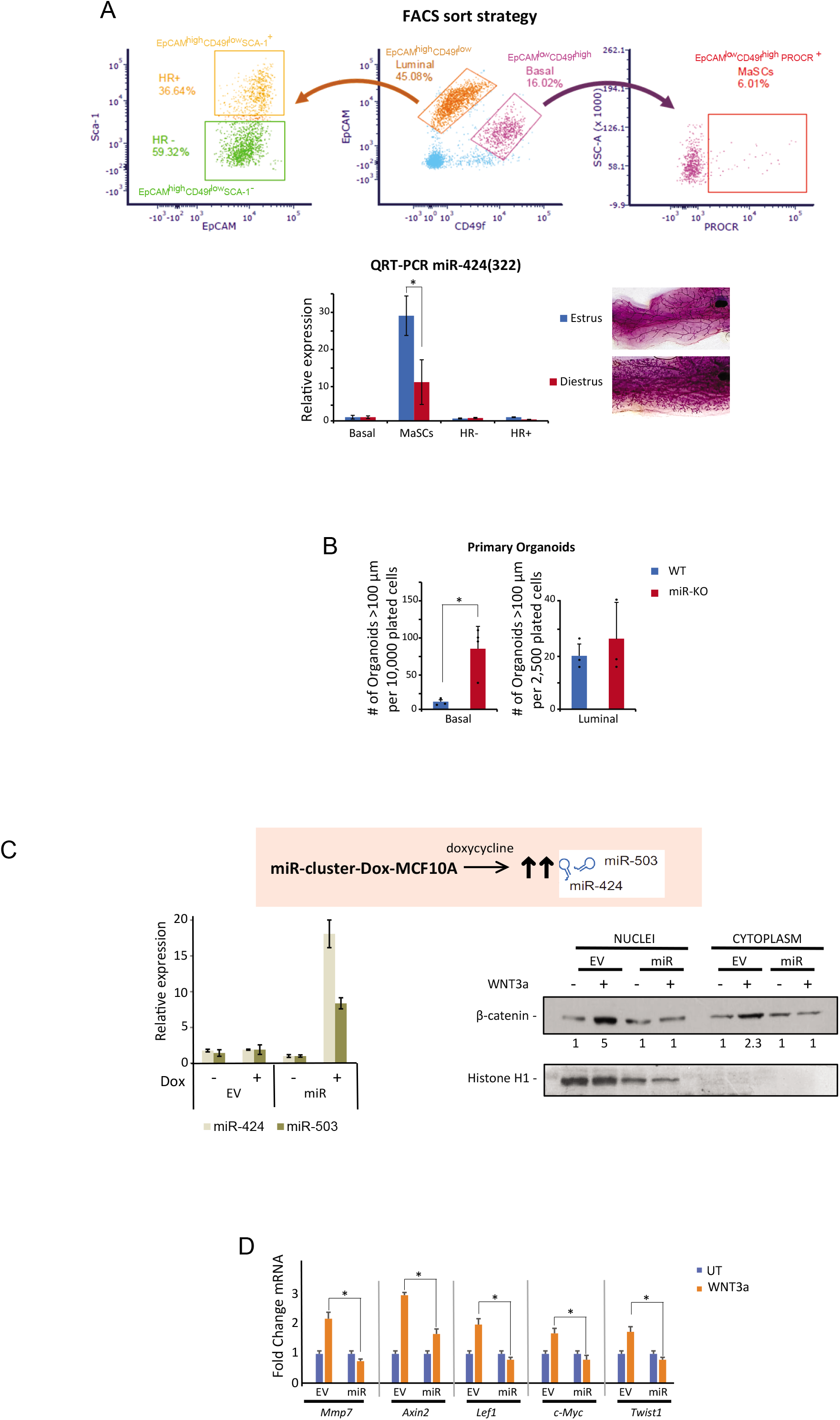
The expression level of miR-424(322)/503 regulates Wnt/β-catenin activity. A) The upper panel indicates the FACS strategy followed to purify the different mammary epithelial subpopulations found in the mouse mammary gland (basal, luminal HR-, luminal HR+, and MaSCs). The lower panel shows the normalized expression of miR-424(503) revealing that MaSCs express the highest levels. FACS was performed at two different stages of the ovarian cycle (estrus and diestrus). Carmine-red staining of the structure of the mammary tree at the time of purification is also provided. B) The bar graphs indicate the number of primary organoids formed when FACS sorted WT or miR-424(322)/503 mammary epithelial basal and luminal cells from female mice at diestrus (4-6 months old) were cultured as organoid. C) The western blot shows the difference in nuclear accumulation of β-catenin when dox-induced (100ng/ml for 3 days) miR-cluster-Dox-MCF-10A cells (miR) and miR-control-Dox-MCF-10A cells (EV, empty vector) were exposed to of WNT3a ligand (100ng/ml for 6 hours). The numbers below the blots indicate the quantitation by densitometry of protein expression relative to EV without WNT3A. Histone H1 staining is shown to control for nuclear-cytosolic purification. The upregulation of miR-424 and miR-503 when dox was added to the media was quantified by QRT-PCR and is shown in the bar graph. D) The panel shows the lack of induction of bona fide WNT targets in miR-over expressing cells from panel C compared with EV control. For all panels, the results shown are the average of n≥3 or representative images of n≥3 independent experiments. The asterisk represents a p-value <0.05.

Based on the high level of expression of miR-424(322)/503 in MaSCs and its association with Wnt-signaling we reasoned that its loss may impact the number of MaSCs in the mammary gland. Thus, we compared the amount of MaSCs present in WT and KO animals. FACS-based strategies to obtain MaSCs only render enriched populations. Thus, the number of MaSCs is normally estimated by using functional assays. For this, we performed a mammary organoid assay (Zhang *et al*, 2016). This assay is based on the ability of MaSC and luminal precursors to form colonies when grown in matrigel suspension and functions as a surrogate marker of the number of MaSCs with repopulation ability. Here, we observed that basal-containing MaSCs (EpCAM^low^CD49f^high^) from KO mice that were FACS-purified and grown in this assay consistently formed more organoids than those from age and stage-matched WT cells (Fig. 2B). As expected, the increased ability to form organoids was maintained when primary structures were disaggregated and plated again to form secondary organoids (Supplemental Fig. S2). Finally, the number of luminal precursors forming colonies observed when luminal (EpCAM^high^CD49f^low^) cells were plated did not significantly change between WT and KO cells, indicating that the absence of miR-424(322)/503 is MaSC specific.

Next, we transitioned our studies to a tractable model where miR-cluster expression and WNT/β-catenin activation could be fully controlled, miR-cluster-Dox-MCF-10A (Llobet-Navas *et al*., 2014a; Llobet-Navas *et al*., 2014b). This is an in vitro model where the expression of miR-424/503 can be experimentally increased at will by using a doxycycline (Dox) inducible system in human basal MCF-10A cells. Additionally, these cells accumulate nuclear β-catenin and upregulate Wnt-responsive genes upon the addition of WNT-ligands (Fig. 2C-2D). Notably, upon upregulation of miR-424/503, both the accumulation of β-catenin and the upregulation of Wnt targets induced by adding WNT3a ligand into the culture media was strongly reduced (Fig. 2C-2D). This result confirms that miR-424/503 expression has a role in modulating canonical Wnt-signaling.

Generally, microRNAs exert their function by binding to a 20-22 nucleotide sequence in the 3’UTR of targeted mRNAs reducing their overall expression by multiple mechanisms (Grimson *et al*, 2007). Thus, to explain the impact of miR-424/503 on WNT/β-catenin signaling, we hypothesized that one or more of its targets are components of this pathway. Computational prediction of targets based on seed sequence conservation as well as non-conserved sites is a widely used strategy to identify miRNA targets. We used the TargetScan algorithm (http://www.targetscan.org), which has been shown to have one of the highest specificities in preselecting putative target genes (Shirdel *et al*, 2011). Notably, TargetScan identified several components that positively regulate the canonical Wnt-pathway and that could explain its activation upon miR-424/503 loss. These putative targets include Wnt-ligands, receptors, and proteins associated with signal transduction. In fact, two of these, *LRP6* and *TBL1XRL*, were in the top 10% of the list when the putative targets were ordered based on their probability of conserved targeting score (PCT) (Fig 3A).

**Figure 3.**
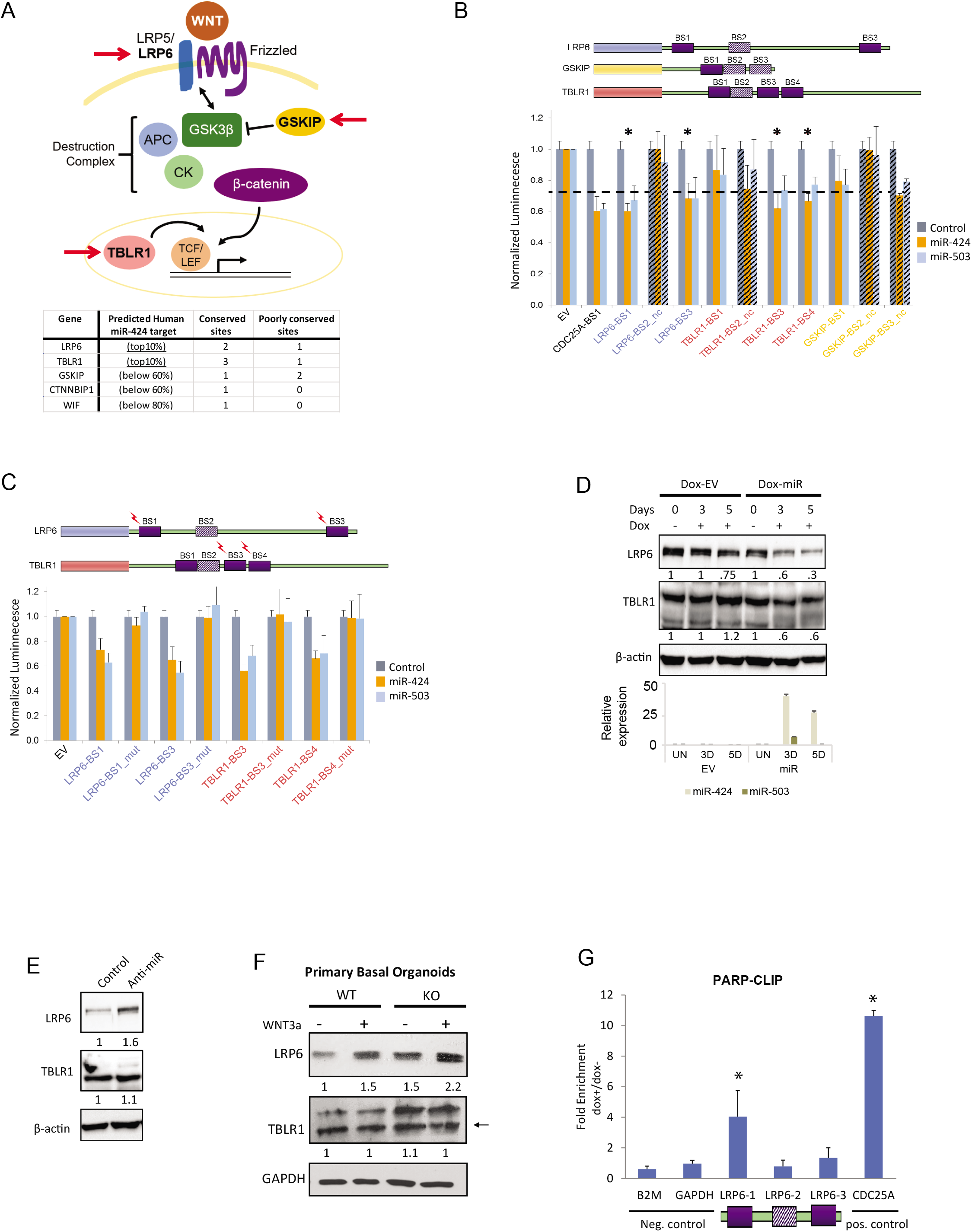
Mir-424/503 cluster targets the Wnt co-receptor LRP6. A) The table shows putative miR-424(322)/503 targets selected from the TargetScan database and ranked based on their probability of conserved targeting. A schematic representation of the canonical Wnt pathway with arrows indicating the putative targets that were investigated is also shown. B) The upper panel illustrates the location of the conserved and non-conserved predicted microRNA binding sites in the 3’UTR of 3 selected target genes. The bar graph compares the normalized expression of a firefly luciferase reporter (luc) plasmid (EV) after transfection in 293T cells and the same luc reporter with a modified 3’UTR containing predicted microRNA binding sites. The values represent the ratio of luc expression when the reporter was co-transfected with synthetic miR-424 and miR-503 vs. a control non-targeting sequence. The bars with a diagonal line pattern indicate non-conserved sites. The asterisk represents a p-value <0.05. A known miR-424/503 binding site located in the 3’UTR of CDC25 is used as a positive control. (C) Luciferase assays showing that the effect of the miR-424 and miR-503 on luc expression is abrogated when the seed sequence in the microRNA binding sites was mutated. The study was performed in four sites statistically significant from panel B. D) Western blot of LRP6 and TBLR1 in the miR-cluster-Dox-MCF-10A model upon upregulation of miR-cluster with 100 ng/ml of doxycycline for the specified number of days. The numbers below the blots indicate the quantitation by densitometry of protein expression relative to β-actin. The expression level of miR-424 and miR-503 is indicated in the bar graph below the blots. E) Western-blots of LRP6 and TBLR1 in parental MCF-10A cells after transfection with miRNA hairpin inhibitors against miR-424/503. F) Western blot of LRP6 and TBLR1 expression in WT and miR-424/503-KO basal cell-derived organoids after 14 days in culture with and without Wnt3a (100ng/ml). (G) Enrichment of the putative LRP6 binding sites bound to AGO2 in miR-cluster-Dox-MCF-10A cells after the miR-424/503 cluster is induced with 100ng/ml of dox for three days. A CDC25A-binding site is included as a positive control and a region of B2M and GAPDH are included as negative controls. For all panels, the results shown are the average of n≥3 or representative images of n≥3 independent experiments. The asterisk represents a p-value <0.05.

As MaSCs cells are sensor cells that react to Wnt-ligands, we decided to focus our studies on signaling components. Thus, we investigated the top two predicted putative targets, *Lrp6* and *TBL1XRL*, which contain 2 and 3 conserved binding sites respectively (Fig 3A and 3B). To investigate whether these putative target genes are regulated by miR-424(322)/503, we first utilized a reporter system measuring the capacity of each predicted binding site to regulate gene expression upon experimental upregulation of miR-424 and miR-503 microRNAs. We then cloned a portion of the gene 3’UTR containing the predicted wild-type binding site downstream of a luciferase reporter gene. Next, we co-transduced 293T cells with the corresponding Luc-UTR reporter, a Renilla luciferase expression vector lacking any UTR and used for normalization purposes, and microRNA mimics corresponding to miR-424(322) and miR-503. Twenty-four hours after transfection, we quantified the expression of the Luc reporter (Fig 3B). These experiments revealed that miR-424 and miR-503 were able to significantly attenuate the expression of Luc when both conserved sites in *LRP6* and 2 out of the 3 conserved sites in *TBL1XR1* were cloned in the 3’UTR. The reduction in Luc expression was equivalent to the reduction observed in the positive control (CDC25A binding site) (Fig. 3B). In contrast, none of the non-conserved sites showed any effect. We also cloned the binding sites, conserved and non-conserved, from *GSKIP*, a predicted target that had a much lower PCT value. Notably, none of its biding sites had any effect on the expression of the reporter (Fig. 3B). To finally demonstrate that the reduction in reporter expression was mediated by the interaction between the miRNA and the targeted mRNA, we mutated the seed sequence of the binding sites (Grimson *et al*., 2007) to disrupt microRNA binding. As expected, this completely abrogated any reduction in reporter expression (Fig 3C).

Next, we tested if the miR-424/503 can modulate the endogenous levels of LRP6 and TBLR1. First, we used the miR-cluster-Dox-MCF-10A model. Here, upon dox induction of the microRNA cluster, western-blot studies showed a clear downregulation in LRP6 protein while a smaller effect was observed for TBLR1 (Fig. 3D). As expected, independent transduction of miR-424 and miR-503 mimics in MCF-10A cells showed that both can downregulate LRP6 (Supplemental Fig. S3). To complement our studies, we also performed loss-of-function in MCF-10A cells. For this, we blocked miRNA-424/503 with the introduction of miRNA hairpin inhibitors into cells. In agreement with the gain of function studies, we observed a clear upregulation of LRP6 while almost no effect was found for TBLR1 (Fig. 3E). Finally, we compared the expression of these putative targets in WT and KO basal-derived organoids in the presence or absence of WNT3a ligand. Again, KO cells showed higher basal expression levels of LRP6 than the WT counterpart, while no main differences were observed for TLBR1 (Fig. 3F). Overall, these results support that miR-424/503 modulates LRP6 expression, but not TLBR1.

MicroRNAs downregulate the expression of targeted genes by recruiting mRNAs to the RNA-induced silencing complex (RISC) (Grimson *et al*., 2007). To finally demonstrate the role of miR-424/503 on LRP6 expression, we performed additional validation using PAR-CLIP (Hafner *et al*, 2010). Briefly, PAR-CLIP is a biochemical assay where the RISC complex and the mRNAs being targeted are immunoprecipitated (IP) using antibodies for the RISC component AGO2. The amount of a particular mRNA bound to RISC is enriched in the IP when a microRNA that modulates its expression is experimentally upregulated. Additionally, the enrichment of RISC in a particular binding site can be determined by simple QRT-PCR. As expected, we observed a significant enrichment of *LRP6* mRNA bound to RISC when the miR-cluster was induced in the miR-cluster-Dox-MCF-10A model (Fig. 3G).

Overall, our study provides compelling evidence that LRP6 is regulated by miR-424(322)/503 in the mammary epithelium and that its loss promotes upregulation of the WNT/β-catenin signaling. As LRP6 is a rate-limiting factor of the WNT/β-catenin pathway (Brennan & Brown, 2004) it is likely that LRP6 is the most significant miR-424(322)/503 target for WNT pathway regulation. We have previously shown that miR-424/503 targets genes such as *IGF1R* and *BCL2* in the mammary epithelium and that its loss affects developmental processes (involution) and tumorigenesis (Llobet-Navas *et al*., 2014a; Rodriguez-Barrueco *et al*., 2017). The newly discovered regulation of WNT-signaling presents a clearer and more complete view of the function of this microRNA-cluster modulating mammary epithelial fate. Thus, to integrate our results with the current understanding of mammary gland development and cancer we propose the following model (Fig. 4). During regular ovarian cycles, at the diestrus phase, there is an increase in the mammary epithelial content of the mammary gland, the purpose of which is to prepare the mammary gland for a potential pregnancy. If pregnancy occurs, gland remodeling becomes widespread (Macias & Hinck, 2012). Expansion of the mammary epithelium during diestrus and pregnancy is mediated by a profound and transient increase of the MaSC pool (Asselin-Labat *et al*., 2010; Joshi *et al*., 2010; van Amerongen *et al*, 2012). This is due to the activation of Wnt/β-catenin signaling in MaSCs responding to RANK and WNT ligands secreted from luminal/HR+ cells (Alexander *et al*., 2012; Goel *et al*., 2012). After expansion, the mammary epithelium involutes to return to its original state; this process includes reduction of the number of MaSCs (Asselin-Labat *et al*., 2010). During periods of epithelial expansion, miR-424(322)/503 is reduced in MaSCs to promote Wnt/β-catenin signaling. During periods of epithelial regression miR-424(322)/503 upregulation in luminal and basal cells attenuates BCL2 and IGF1R expression inducing cell death/apoptosis (Llobet-Navas *et al*., 2014a; Rodriguez-Barrueco *et al*., 2017) and reducing Wnt/β-catenin signaling in MaSCs by targeting LRP6. Loss of miR-424(322)/503 in breast cancers (Rodriguez-Barrueco *et al*., 2017) and miR-KO mice sensitizes MaSCs to WNT stimuli during phases of epithelial expansion and promotes aberrant survival of mammary epithelial cells due to upregulation of BCL2 and IGF1R during periods of epithelial regression. This accumulates through the female lifetime promoting tumorigenesis (Rodriguez-Barrueco *et al*., 2017).

**Figure 4.**
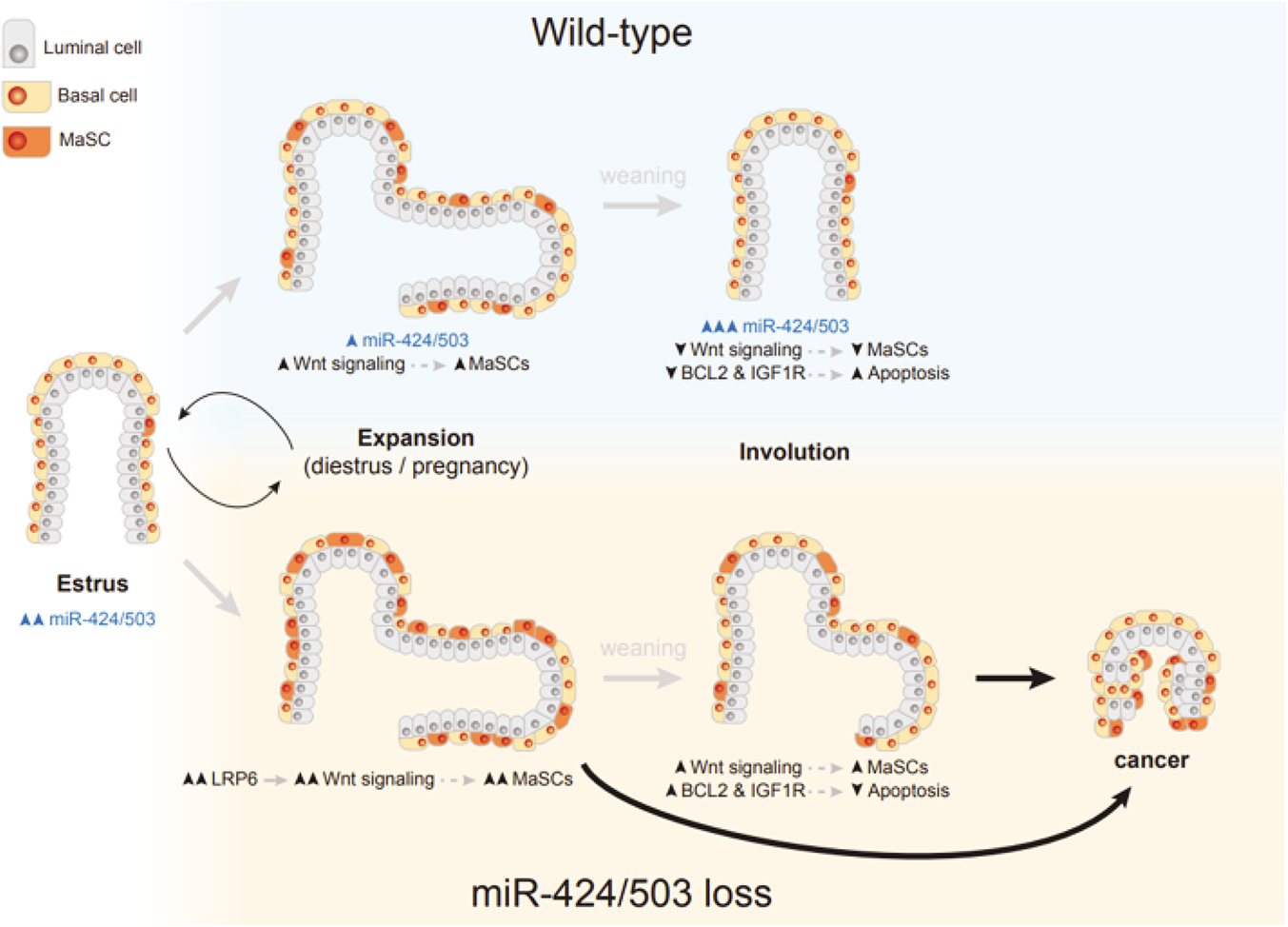
Model of miR-424/503 function in the mammary gland. This graphic illustrates the functional model presented in the discussion for miR-424/503 in the mammary gland and how its loss promotes breast cancer.

Finally, an important finding of our study is the high level of miR-424/503 deletions in TNBCs of the BL2 and M subtypes. It has been long recognized that Wnt-signaling is activated in some breast cancers, especially in the TNBC group (Lehmann *et al*., 2011). However, the molecular alterations linked to it have remained elusive. Thus, our study has unveiled a molecular alteration that can explain a large fraction (∼25%) of all cases.

In summary, the results presented here unveil an unknown link between the miR-424/503, the regulation of WNT-signaling, MaSC fate, and tumorigenesis.

## Author Contributions

JS, and DLL conceived, designed, and supervised the research and wrote the manuscript. JS, DLL, EN, and RRB developed the experimental methodology. JY and KKY coordinated and performed the computational analysis. EN and RRB had equal contributions coordinating and performing the wet lab studies described in the text. RLW and PM contributed to the animal studies. All authors contribute to data analysis and manuscript editing and wrote different parts of the material and method section.

## Acknowledgments

This study has been partially funded by the NY-state breast cancer grant program (Peter T. Roley) (JS); by the Instituto de Salud Carlos III (ISCIII) through the projects CP17/00063 (cofounded by European Regional Development Fund (ERDF) “a way to build Europe”) and MS17/00063 (cofounded by the European Social Fund (ESF), “investing in your future”), and by the Ministerio de Ciencia, Innovación y Universidades Gobierno de España, through the projects RyC-2016-19671 and RTI2018-095611-A-I00 (DLL). Ministerio de Ciencia, Innovación y Universidades Gobierno de España, through the projects RyC-2016-19671 and RTI2018-095611-A-I00 (RRB). The National Cancer Institute (NCI) of the National Institutes of Health (NIH) under award number F31CA232691 (EN) The data in this paper were used in a dissertation as partial fulfillment of the requirements for a Ph.D. degree at the Graduate School of Biomedical Sciences at Mount Sinai.

## Supplemental Figures

**Supplemental Figure 1.**
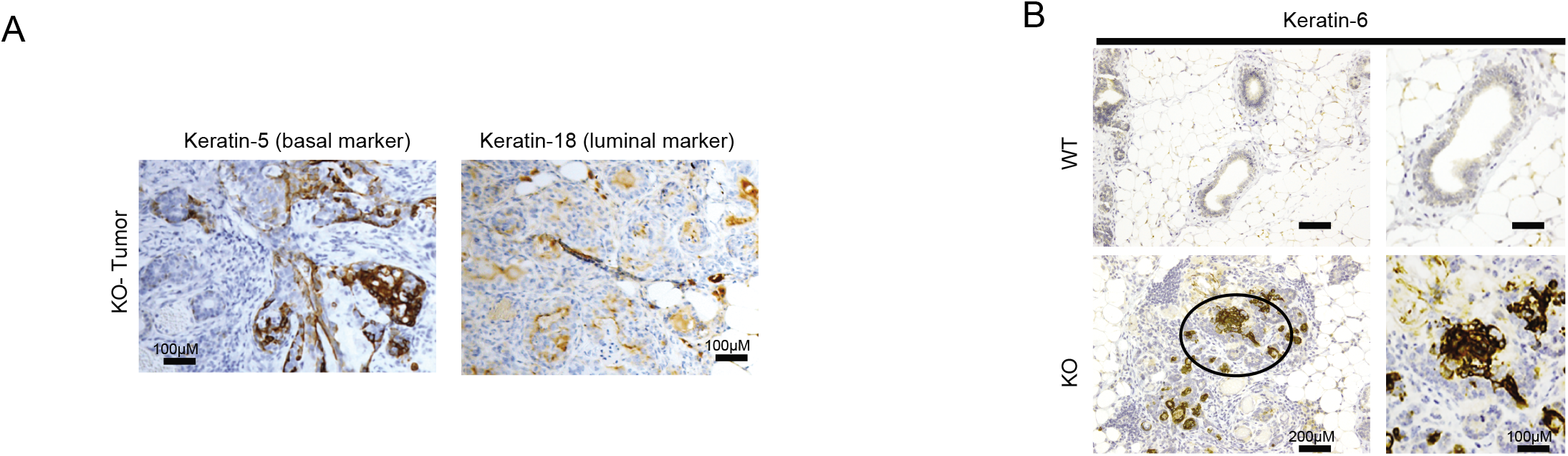
Activation of Wnt signaling in breast cancers. A) Tumors emerging in one-year-old miR-424(322)/503-/- female mice present marker of basal (keratin-5) and luminal (keratin-18) cells. B) Immunohistochemistry reveals that keratin-6 accumulates in tumors emerging in female miR-424(322)/503^-/-^ mice. All results shown are representative images of n≥5 independent experiments.

**Supplemental Figure 2.**
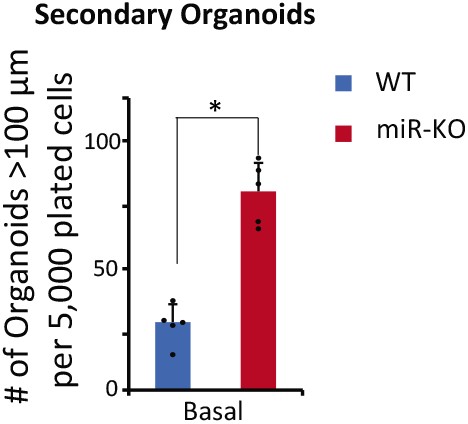
MiR-424(322)/503-KO basal cells show enhanced organoid forming potential. The bar graphics indicate the number of secondary organoids formed when FAC-sorted WT or miR-424(322)/503-KO mammary epithelial basal cells from female mice at diestrus (4-6 months old) were growing in organoid forming cultures. The results shown are the average of n≥3 independent experiments. The asterisk represents a p-value <0.05.

**Supplemental Figure 3.**
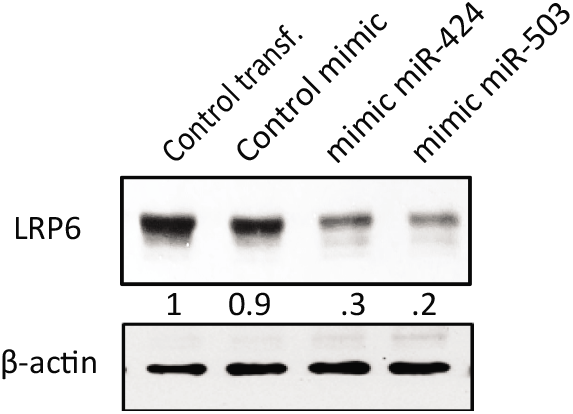
MiR-424(322)/503 targets the WNT co-receptor LRP6. WT-blot showing reduction expression of endogenous LRP6 72 hours after MCF-10A cells were transduced with miRNA mimics (100nM). The numbers below the blots indicate the quantitation by densitometry of protein expression relative to β-actin.

## Materials and Methods

### Animal models

The generation of miR-424(322)/503-/- animals has been previously described (Llobet-Navas *et al*., 2014a; Rodriguez-Barrueco *et al*., 2017).

### Patient data analysis

Patient data from TCGA-BRCA (Cancer Genome Atlas, 2012) consists of breast cancer samples with available whole-genome DNA copy number alterations, mRNA expression data, clinicopathological and reverse-phase protein array (RPPA) data. Patient data from METABRIC (Curtis *et al*., 2012) was analyzed for whole-genome DNA copy number alterations and mRNA expression data.

### Cell culture

Cell lines were obtained from the American Type Culture Collection (ATCC). MCF10A cells were cultured in DMEM/Ham’s F-12 media supplemented with 1% Penicillin-Streptomycin, EGF (20 ng/ml), insulin (10 µg/ml), cholera toxin (100 ng/ml), hydrocortisone (500 ng/ml) and 5% horse serum. We cultured 293T cells in DMEM media with 1% Penicillin-Streptomycin and 10% FBS. In some cases, described in the text, Wnt3a ligand (R&D systems 5036-WN) at 100ng/ml was added to the culture. MiRNA inhibition in MCF-10A cells was achieved by transducing cells with a miRNA inhibitor against hsa-miR-424-5p and hsa-miR-503-5p (Genecopoeia #HmiR-AN0494-AM03 and #HmiR-AN0550-AM03 respectively). Control cells were generated by transducing a scrambled control (Genecopoeia #CMIR-AN0001-AM039).

### miR-cluster-Dox-MCF-10A

The generation of these cells has been previously described (Llobet-Navas *et al*., 2014a; Rodriguez-Barrueco *et al*., 2017). Doxycycline was added to the media at 100ng/ml to induce the expression of miR-424/503.

### RNA and protein analysis

To perform microRNA expression analysis by qRT-PCR, RNA was extracted using the miRVANA miRNA isolation Kit (Ambion #AM1561) according to manufacturer’s instructions. Reverse transcription and qPCR were performed according to the TaqMan microRNA Assay protocol and as previously described using probes for murine miR-322, miR-503, and snoRNA-202 (ThermoFisher #001076, 002456, and 001232, respectively). The thermal cycler conditions were: activation at 95°C for 10 minutes, denaturation 95°C for 15 seconds, and annealing/extension 60°C for 1 minute (repeat for 40 cycles). Ct values (2-ΔΔCT) were calculated by normalizing to an endogenous reference gene (snoRNA-202). To assess the expression of Wnt/ β-catenin target genes, total RNA was converted into cDNA using the High Capacity cDNA Reverse Transcription Kit (Roche #4368814) according to manufacturer’s instructions. Two microliters of the reverse transcription reaction were used as a template for real-time detection on a BioRad MYIQ Single-Color Real-Time PCR Detection System. For coding genes, the Real-time PCR reaction was performed with 10uL FastStart SYBR Green Master Mix (Roche #04673492001), 2 uL of complementary DNA (cDNA), and 1uL of each primer in RNAse-free water adjusted to 20uL volume reaction. The following were the thermal cycler conditions: AmpliTaq activation 95°C for 3 minutes, denaturation 95°C for 10 seconds, and annealing/extension 60°C for 30 seconds (repeat 40 times). Triplicate Ct values were further analyzed (2-ΔΔCT) by normalizing to an endogenous reference gene (GAPDH). Results are presented as the relative mRNA amount compared to the untreated samples. Primers used: *GAPDH* (F CATCTTCTTTTGCGTCGC, R AAAAGCAGCCCTGGTGAC), *MMP7* (F GAGTGAGCTACAGTGGGAACA, R CTATGACGCGGGAGTTTAACAT), *AXIN2* (F CAACACCAGGCGGAACGAA, GCCCAATAAGGAGTGTAAGGACT), *LEF1* (F AGAACACCCCGATGACGGA, R GGCATCATTATGTACCCGGAAT), *MYC* (F TCCCTCCACTCGGAAGGAC, CTGGTGCATTTTCGGTTGTTG), *TWIST1* (F GTCCGCAGTCTTACGAGGAG, R GCTTGAGGGTCTGAATCTTGCT).

### Western blotting

Cells were lysed in RIPA buffer with 30 mM NaF, 1 mM Na3VO4, 40 mM β-glycerophosphate, 0.1 mM PMSF, and protease inhibitors. Protein concentrations were determined by the Protein Assay Kit (Bio-Rad #500-0006). We subjected equal amounts of proteins to SDS-PAGE and transferred them to nitrocellulose membranes (GE Healthcare #10401197). Non-specific binding was blocked by incubation with TBST plus 5% non-fat milk. We incubated the membranes with primary antibodies overnight at 4°C and with secondary HRP-conjugated antibodies (Amersham #NA9350V, #NA931V, and #NA934V) for 1 hour at room temperature. Signal was detected with SuperSignal West Pico or Dura Extended Duration Chemiluminescent Substrate (Thermo #34079 or #34076). The primary antibodies we used for these studies are: anti-β-catenin (abcam #ab2365), anti-LRP6 (Cell Signaling #3395), anti-TBLR1 (Bethyl #A300-408A), and anti-GAPDH (Cell Signaling #14C10). Quantification of western blot signal was performed using a ChemiDoc Imaging Systems and the Image Lab Software package (Bio-Rad).

### Whole Mount Preparations and Immunohistochemistry

Mammary glands were fixed in formalin (Fischer #175) for immunohistochemistry (IHC) analysis. Whole-mount carmine red staining to visualize the mammary tree branching was prepared as follows: mammary glands were dissected and fixed in paraformaldehyde 4% for 1 hour. Afterward, samples were stained with carmine red (Sigma #C1022) overnight at room temperature. The next day samples were dehydrated in increasing concentrations of ethanol (70, 95, and 100%) and ultimately fixed in xylene.

Formalin-fixed paraffin-embedded samples were first heated at 100°C for 3 minutes on a heat block to melt the paraffin. Subsequently, samples were deparaffinized by serial incubations with xylene for 3 minutes, 100% EtOH for 3 minutes, 95% EtOH for 3 minutes, and distilled water for 2 minutes. Peroxidase inactivation and antigen retrieval were achieved by incubating samples in 1% H2O2 for 15 minutes at room temperature and incubating slides with citric buffer (2mM citric acid, 8mM sodium citrate) in a steamer for 30 minutes. Samples were washed twice in PBS for 5 minutes and incubated in 10% whole goat blocking serum diluted in 2% BSA-PBS for 30 minutes at room temperature. Samples were then incubated in primary antibody diluted in 2% BSA-PBS+0.01% sodium azide for 2 hours at room temperature. Antibodies were used at 1:300 Anti-Cytokeratin 18 antibody (Abcam #ab668), 1:1000 Anti-Cytokeratin 5 (Covance #PRB-160P). The remaining antibodies used were the same as for WT-blotting but used at 1:100. Samples were then washed in PBS and incubated in 1:500 biotinylated anti-Rabbit IgG made in goat diluted in 2% BSA-PBS for 30 minutes at room temperature. Afterward, samples were washed and exposed to peroxidase substrate (Vector Laboratories #PK-6100) for 30 minutes at room temperature and subsequently permeabilized with PBS-0.5% Triton. Thereafter samples were incubated in chromogen 3,3’ Diaminobenzidine (DAB) and then washed in distilled water and counterstained. Counterstaining was performed by treating samples with hematoxylin for 1 second, dipped in 1% Hydrochloric acid, and finally washed in ammonia water for 1 second. Finally, dehydration was performed by incubating samples in 95% EtOH for 2 minutes, 100% EtOH for 2 minutes, and xylene for 4-5 minutes and ultimately mounted with a coverslip.

### Nuclear fractionation

NF was performed using the NE-PER™ Nuclear and Cytoplasmic Extraction Reagents and following the instructions provided by the vendor (Life Technologies # 78833).

### Luciferase reporter assays

To measure the targeting activity of miR-424(322)/503, we cloned oligonucleotides corresponding to the region of the 3’UTR containing each miRNA binding site (plus 30bp upstream and downstream) downstream of the luciferase reporter in the pMIR-REPORT vector (Life Technologies #AM5795M) using restriction enzymes. For the mutagenesis, the same method was followed except that we randomly mutated the 8 base pair sequence corresponding to the miRNA seed sequence. To measure luciferase activity, 293T cells were seeded at 70% confluence in 96-well plates. 24 hours later, we co-transfected the cells with 50 ng of pMIR-REPORT constructs containing the 3’UTR sequences together with 50 ng of a Renilla housekeeping control plasmid and 100 nM of miR-424 or miR-503 mimics or a miRNA negative control (Dharmacon #C-300717-05, #C-300841-05, or #CN-001000-01-05, respectively) using the TransIT-X2 reagent (Mirus 6003). 24 hours after transfection, we measured the relative luciferase units (RLU) using the Dual-Glo Luciferase Assay System (Promega #E2949).

### PAR-CLIP

The PAR-CLIP assay to measure the enrichment of miR-424(322)/503 mRNA targets bound to AGO2 was performed as described previously (Hafner *et al*., 2010). Briefly, cells were pretreated with 50 uM of 4-thiouridine (Sigma #T4509) overnight and crosslinked at 150mJ/cm2 at 365nm UV on ice. Cells were reconstituted in lysis buffer [2.5 mM Hepes pH 7, 50 mM NaCl, 10% glycerol, 1% Triton X-100, proteinase inhibitor (Roche #04693159001), 0.2 mM DTT and 1 U/µl RNAseOUT (Invitrogen #10777-019)]. Mild (5 U/µl) RNase-T1 (Fermentas #EN0541) digestion was performed at 22°C for 15 minutes. Immunoprecipitation was performed using A-beads (Roche #11719408001), G-beads (Roche #11719416001), and 10 µg of anti-AGO2 antibody (Abnova #H00027161-M01) overnight at 4°C. Samples were subsequently washed twice with washing buffer 1 (50 mM Tris pH 7.5, 150 mM NaCl, 0.1% NP-40 and 1 mM EDTA) at 4°C for 30 minutes, digested with RNase-T1 (20 U/µl) at 22°C for 15 minutes, washed once with washing buffer 2 (50 mM Tris pH 7.5, 500 mM NaCl and 0.1% NP-40) at 4°C for 30 minutes and finally washed twice with washing buffer 3 (50 mM Tris pH 7.5 and 500 mM NP-40) at 4°C for 30 minutes. Samples were centrifuged for 5 minutes at 5000rpm at 4°C and supernatant treated with 5 mg/ml of proteinase K (New England Biolabs #P8102) for 1 hour at 50°C. Finally, the extract was lysed, and RNA extracted using the miRVANA miRNA isolation Kit (Ambion #AM1561) according to the instructions provided. Primer sequence used to measure LRP6 binding site (BS) enrichment in RISC by qRT-PCR: LRP6-BS1 (F TTTGTACAGAAGAAAAGGAT, R AGTTTGCAAAAATAAAACTT), LRP6-BS2 (F GAATAATGGAAGCCTCTTT, R CTAGAATCATTCCACAGGT) and LRP6-BS3 (F TACCAAGAAGATTAAACTGG, R AGATTAAAGCTTAAGGGAAA). Primers for negative/positive controls were: GAPDH (F CATCTTCTTTTGCGTCGC, R AAAAGCAGCCCTGGTGAC), B2M (F TGCTGTCTCCATGTTTGATGTATCT, R TCTCTGCTCCCCACCTCTAAGT) and CDC25A (F GCCATTCTAGGTAGGGTTTT, R CCTAGCTTTCTGTCCGATAA).

### Mammary gland dissociation and FACS

The 3rd and 4th mammary glands from female mice of the specified age, genotype, and estrous cycle phase were pooled in PBS and minced followed by enzymatic digestion in DMEM/F12 media containing 2 mg/ml Collagenase A (Roche #11088793001) and 100 U/ml Hyaluronidase (Sigma #H3506) for 2 hours at 37°C. Single-cell suspensions were obtained via sequential incubations in pre-warmed 0.25% Trypsin-EDTA followed by 5 mg/ml Dispase II (Life Technologies #17105-041) in PBS with 0.1 mg/ml DNase I (Stemcell #07900), pipetting up and down at each step. Finally, cells were incubated in Red Blood Cell Lysing Buffer (Sigma #R7757) and filtered through a 40 µm cell strainer. Then epithelial cells were obtained using the EasySep Mouse Epithelial Cell Enrichment Kit II (Stemcell #19758) according to the protocol provided. The epithelial cell-enriched suspensions were incubated with the following antibodies: EpCAM-PerCP/Cy5.5, CD49f-APC, PROCR-PE, and Sca-1-BV421 (BioLegend #118220, #313616, #141504, and BD #562729, respectively) to sort the different epithelial subpopulations. To determine cell viability, we used the LIVE/DEAD™ Fixable Aqua Dead Cell Stain (ThermoFisher #L34957). Antibody incubations were performed for 15 minutes on ice before sorting the cells using a FACSAria II cell sorter (BD).

### Organoid culture

FACS sorted cells were seeded into 24-well ultra-low attachment plates at a density of 5,000 or 10,000 cells/well in EpiCult-B Mouse Medium (StemCell #05610) supplemented with 5% FBS, 10 ng/ml EGF, 20 ng/ml bFGF, 4 μg/ml heparin, 5 μM Y-27632, and 5% Matrigel (Corning #354230). After 7 days in culture, the organoids were measured. For the generation of secondary organoids, the organoids were dissociated by incubating primary organoids in 0.25% Trypsin-EDTA in PBS for 5 minutes at 37°C followed by mechanical disruption with a P1000 pipette. Finally, cells were filtered using a 40 µm cell strainer to ensure a single-cell suspension and seeded them again into 24-well ultra-low attachment plates and counted the number of organoids >100 μm in diameter after 7 days as described above. In some cases, described in the text, Wnt3a ligand (R&D systems 5036-WN) at 100ng/ml was added to the culture.

